# Source to sink partitioning is altered by the expression of the transcription factor AtHB5

**DOI:** 10.1101/2022.06.15.496323

**Authors:** L Raminger, VN Miguel, C. Zapata, RL Chan, JV Cabello

## Abstract

Carbohydrates are transported from source to sink tissues. The efficiency of such transport determines plant growth and development. The process is finely regulated, and transcription factors are crucial in such modulation. AtHB5 is a homeodomain-leucine zipper I transcription factor, repressed during stem secondary growth. However, its function in this developmental event was unknown. Here, we investigated the expression pattern and role of AtHB5. *AtHB5* localized in conductive tissues: roots, hypocotyls, stems, pedicels, and central leaf veins. Mutant plants exhibited wider and more lignified stems than controls, whereas overexpressors showed the opposite phenotype. Cross-sections of *athb5* mutant stems showed enlarged vascular bundle, xylem, phloem, and petiole areas, whereas *AtHB5* overexpressors exhibited callose deposits. Several genes involved in starch biosynthesis and degradation had altered transcript levels in *athb5* mutants and *AtHB5* overexpressors. Rosette and stem biomasses were enhanced in *athb5* mutants, positively impacting seed yield and lipid content. Moreover, these effects were more evident in debranched plants. Finally, the transport to roots significantly slowed down in *AtHB5* overexpressors.

Altogether, the results indicated that AtHB5 is a negative modulator of sucrose transport from source to sink tissues, and its overexpression diminished plant biomass and seed yield.

**Highlight:** The homeodomain-leucine zipper transcription factor AtHB5 is expressed in different tissues along the life cycle, repressing carbohydrate transport from source to sink and promoting callose and lignin deposition. AtHB5 mutants exhibit physiological differences with the wild-type, impacting seed yield and lipid content.

## INTRODUCTION

Plants keep their energy reserve mainly as carbohydrates. However, storage is not the sole function of these biomolecules; they have other roles like acting as building blocks and osmolytes, required for the antifreezing response, among others (Nägele *et al*., 2010). Carbohydrates are the photosynthesis products in the leaf mesophyll cells. Sucrose is the main transported sugar; it provides energy for the growth and development of roots, flowers, fruits, and seeds (Baker *et al*., 2012; Braun, 2012). It diffuses cell to cell through plasmodesmata and reaches the parenchyma cells of the phloem (Slewinski and Braun, 2010).

Sucrose is carried by SWEET carriers, from the phloem to the apoplast, where it is loaded into the phloem sieve element-companion cell complexes by SUC transporters (Braun and Slewinski, 2009; Ayre, 2011; Chen *et al*., 2012). Sucrose accumulation produces a pressure gradient generating a flow, finishing in sink tissues, where it is stored or used (Lalonde *et al*., 2004; Baker *et al*., 2012). For long-distance sucrose transport, sieve elements are aligned, forming sieve tubes (Evert, 1982). The regulation of carbon partitioning is crucial to achieving grain filling and, ultimately, seed yield (Gifford *et al*. 1984). From the fixed carbon, up to 80 % is exported to sinks (Kalt-Torres and Huber, 1987). Major sinks during the vegetative stage are roots and young leaves, whereas fruits and seeds become dominant during the reproductive one (Wardlaw, 1990). However, the roots are still energy-consuming at the end of the life cycle because they need the energy to provide water and minerals to complete development.

Starch is the most abundant storage form. It accumulates during the day in leaves using a fraction of the carbon assimilated through photosynthesis. During the night, it is consumed, producing sucrose (Smith and Stitt, 2007; Stitt and Zeeman, 2012). Mutant plants that fail to synthesize or degrade starch exhibit reduced growth rates under most conditions (Usadel *et al*., 2008; Yazdanbakhsh and Fisahn, 2011).

Callose is a β-1,3-linked glycan that produces deposits that block cell to cell movement of diverse metabolites. The control of such deposition is crucial for phloem sap transport and resource allocation, impacting plant development (van Bel, 2003; Chen and Kim, 2009; Barratt *et al*., 2011; Xie *et al*., 2011; Piršelová and Matušíková, 2013). The synthesis and degradation of carbohydrates are regulated processes involving many enzymes. Among them, SS1 (SUCROSE SYNTHASE 1) participates in amylopectin synthesis, SS4 (SUCROSE SYNTHASE 4) promotes starch granule initiation, and GBSS is a granule-bound starch synthase (Szydlowski *et al*., 2011; Abt *et al*., 2020). ADG1 and ADG2 are the small and large subunits of ADP-glucose pyrophosphorylase, which catalyzes the rate-limiting step in starch biosynthesis. PGM (PHOSPHOGLUCOMUTASE) catalyzes the reversible conversion of glucose 6-phosphate to glucose 1-phosphate.

At night, starch degradation begins with the phosphorylation of glucan chains by GWD (GLUCAN WATER DIKINASE) and PWD (PHOSPHOGLUCAN WATER DIKINASE) (Ritte *et al*., 2006). This phosphorylation is thought to disrupt the structure of the outer chains, facilitating hydration and subsequent hydrolysis by β-amylases (BAM) proteins (Fulton *et al*., 2008). In Arabidopsis guard cells, the major starch-degrading enzyme is BAM1. When the day starts, BAM1 rapidly mobilizes starch with the chloroplastic AMY3, an ENDOAMYLASE that hydrolyzes α-1,4 bonds within glucan chains (Horrer *et al*., 2016). BAM4 encodes a catalytically inactive protein located in plastids, where it may play a role in regulating starch metabolism (Fulton *et al*., 2008; Li *et al*., 2009).

Genes involved in carbohydrate synthesis and degradation, as well as those encoding sucrose transporters, wall synthesis, determination of stem thickness, and other events crucial for carbon partitioning, are modulated by environmental and illumination conditions (Ruan *et al*., 2014). The regulation occurs at different levels, and the transcriptional one is particularly important.

Stems are the main carbohydrate transport organs, and their morphological characteristics, such as width and height, are related to seed yield (Cabello and Chan, 2019). Studying the molecular mechanisms involved in stem secondary growth, Arabidopsis plants were treated by applying weight, and differential transcriptome analysis was carried out (Ko *et al*., 2004). This analysis resulted in 700 differentially expressed genes, including transcription factors (TF) belonging to several families. Among them, a group of homeodomain-leucine zipper TF appeared as modulated during stem secondary growth (Ko *et al*., 2004). Interestingly, nine were induced, and only one, *AtHB5*, was repressed. AtHB5 can heterodimerize with other specific members of the HD-Zip I subfamily, including AtHB6, AtHB16, AtHB7, and AtHB12 (Johannesson *et al*., 2001). Moreover, a yeast one-hybrid assay revealed that AtHB5 binds the BDL promoter region repressing its expression (De Smet *et al*., 2013). Considering the expression pattern of *AtHB5*, it was localized in the hypocotyl of germinating seedlings and later in the petioles of cotyledons and developing leaves. Interestingly, in adult plant tissues, *AtHB5* was detected in the differentiation zone of main and lateral roots, the vascular tissue of cauline leaves and stems, and pedicels of flowers and siliques (Johannesson *et al*., 2003). Its overexpression in Arabidopsis caused the enhancement of ABA sensitiveness during seed germination and seedling growth (Stamm *et al*., 2017).

Although *AtHB5* was expressed in stems and other conductive tissues, its function in these organs was unknown. In this work, we investigated the role of this TF in carbon partitioning. Studying mutant and overexpressor plants, we determined that AtHB5 is a negative regulator of sucrose transport, affecting carbohydrate partitioning from source to sink tissues and impacting seed yield.

## RESULTS

### AtHB5 negatively modulates lignin deposition impacting secondary growth

Stem maturation involves the up and down-regulation of many genes. Among them, *AtHB5* was the only HD-Zip I transcription factor down-regulated during this event (Ko *et al*., 2004).

To characterize the putative function of AtHB5, we obtained three independent overexpressor lines (*35S:AtHB5*, thereafter named AT5) exhibiting varied expression levels (Supplementary Figure S1) and three homozygous mutant lines (*athb5-1, athb5-2*, and *athb5-3*). AT5, *athb5*, and WT plants did not show significant differences in the main stem height along the life cycle (Figure 1A and Supplementary Figure S2). Regarding stem width, *athb5* mutants were wider than controls, whereas overexpressors showed the opposite phenotype (Figure 1A and Supplementary Figure S2).

**Figure 1.**
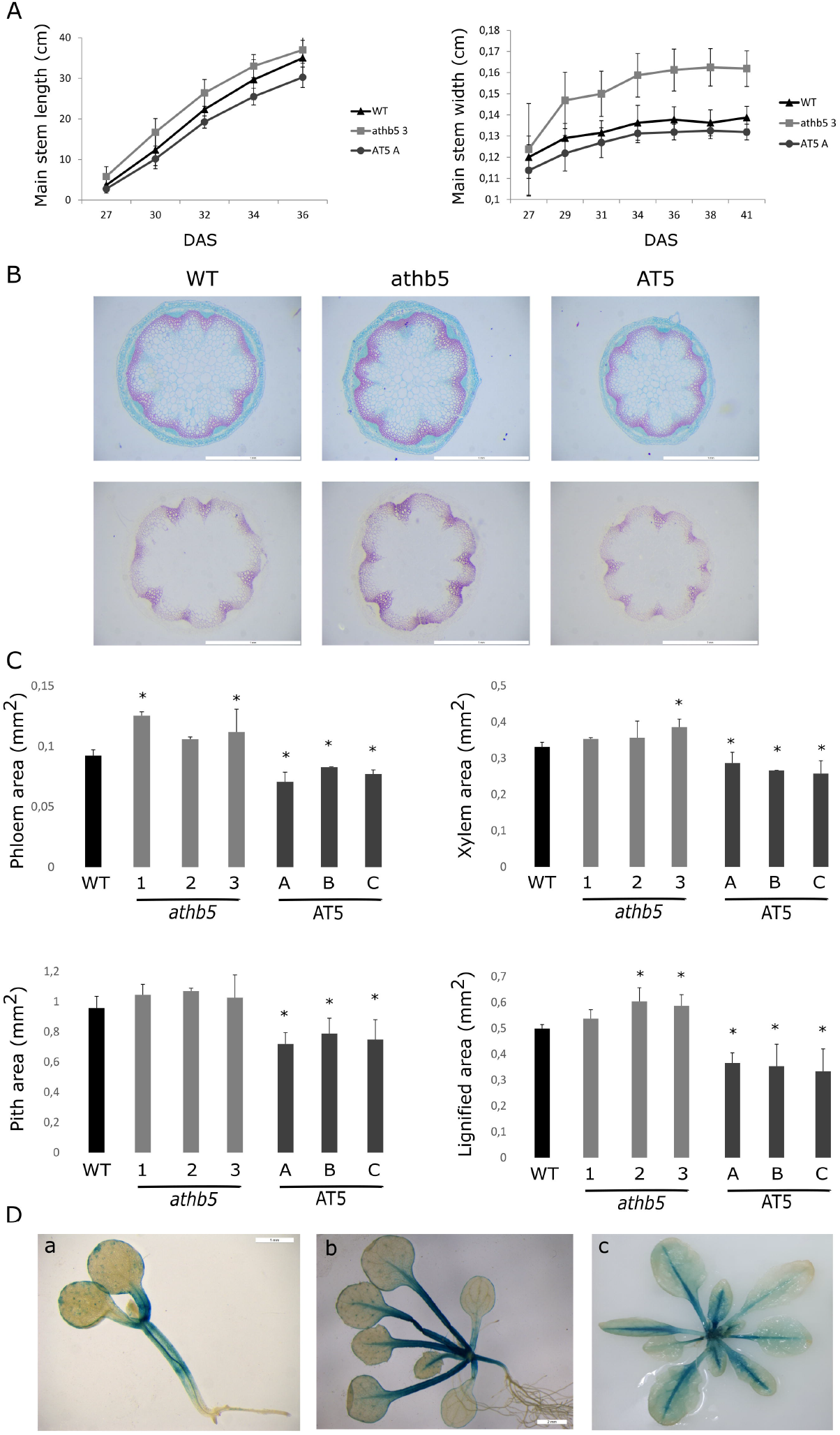
Plants with altered levels of *AtHB5* exhibit stem differential traits. (A) Length and width of the main stem of *athb5* mutant, AT5 overexpressor, and wild-type (WT) plants. (B) Cross-sections were taken from the first stem internode of 10 cm plants stained with Safranin-Fast green solution (upper panel) to visualize xylem and phloem tissues and HCl-Fluoroglucinol (lower panel) to detect lignin deposition. White bars indicate 1mm (C) Quantification of phloem, xylem, pith, and lignified area (mm^2^) using ImageJ software. (D) Histochemical detection of GUS enzymatic activity in *ProAtHB5:GUS* 7-day old (a), 14-day old (b) plants grown in MS, and rosette of 25-day-old plants (c) grown in soil. The white scale bar represents 1 mm in (a) and 2 mm in (b). Quantification of the area in histological sections was performed with four biological replicates per genotype. Length and width measurements were repeated three times with 10 plants per genotype. Bars represent SEM. Differences were considered significant and indicated with asterisks when the P-value was <0.05 (Student’s t-test).

Given that lignin deposition on xylem vessels is one of the main characteristics of secondary growth, this trait was analyzed doing transversal sections stained with Safranin-Fast green or Floroglucinol-HCl. This evaluation indicated less lignin content in the overexpressors, and the contrary in *athb5* mutants (Figure 1B). Moreover, quantification of tissue parameters on stem cross-sections indicated that phloem, xylem, pith, and lignified xylem area showed an increase in mutants and a decrease in overexpressors, in agreement with the staining (Figure 1C).

To better understand the putative role of AtHB5 in stem development, we cloned an 1531 bp segment upstream of the start codon directing the expression of the *GUS* reporter gene (*PrAtHB5:GUS*) and then transformed Arabidopsis plants with this construct. The expression pattern of *AtHB5* was analyzed by histochemistry at different developmental stages. In 7-day-old seedlings, hypocotyls and petioles were stained, and this expression continued until day 14, adding the vascular tissue of leaves. The expression remained active in petioles and vascular tissue of 21-day-old plants (Figure 1D).

Altogether, these results indicated that *AtHB5* is expressed in conductive tissues at least in early developmental stages and negatively modulates lignin deposition and stem width enlargement.

### Altered AtHB5 levels impact stem anatomy and carbohydrate transport

To further characterize the differential stem phenotype dependent on *AtHB5* levels, glucose, sucrose, and starch concentrations were determined in rosette leaves throughout the day cycle. Interestingly, glucose content was higher in mutants ending the day (at 6 pm), whereas sucrose showed elevated levels at 10 am, both compared to WT (Figure 2A). Starch accumulated in AT5 plants towards the end of the light cycle (10 pm), while the opposite happened in *athb5* mutants (Figure 2A). These observations indicated significant differences in carbohydrate transport to sink tissues depending on *AtHB5* levels (Supplementary Figure S3).

**Figure 2.**
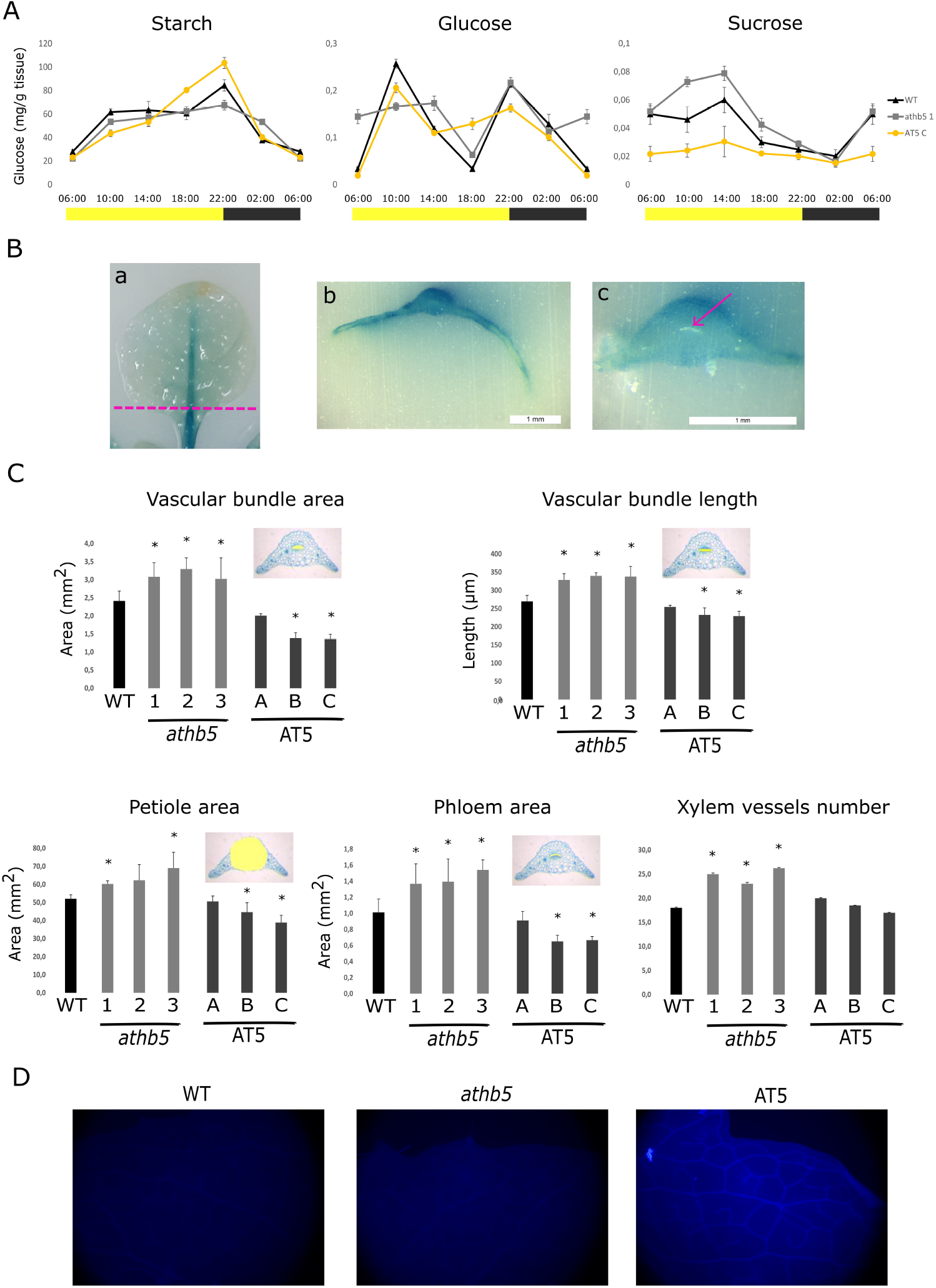
Carbohydrate contents, petiole morphology, and callose deposition indicate a deficient sucrose transport in AT5 plants. (A) Starch, glucose, and sucrose contents in 25-day-old rosette leaves from *athb5* mutant, AT5 overexpressor, and WT plants grown in normal growth conditions. Samples were taken during the day at the hours indicated on the x-axis; grey bars signal indicate the night period and yellow bars the day. (B) Histochemical detection of GUS enzymatic activity in *ProAtHB5:GUS* rosette leaves (a), histological cuts from a rosette leaf (b), and a petiole (c). White bars indicate 1 mm and the pink arrow shows the lack of GUS activity in the phloem tissue. (C) Vascular bundle area, vascular bundle length, petiole area, phloem area, and xylem vessel number were quantified in petiole cross-sections using ImageJ. Carbohydrate determination was repeated twice with four biological replicates randomized from eight individual plants per genotype. Quantification of the area in histological sections was performed with four biological replicates per genotype. Bars represent SEM. Differences were considered significant and indicated with asterisks when the P-value was <0.05 (Student’s t-test). (D) WT, *athb5*, and *AT5* rosette leaves stained with aniline blue detecting callose with a Fluorescence microscope.

Given that GUS staining was detected in the vascular tissue of petioles and leaves, we performed histological cuts of *ProAtHB5:GUS* petioles. Notably, the whole petioles were stained, except the region corresponding to phloem tissue (Figure 2B). To know if *AtHB5* expression in the petioles causes anatomical changes, we performed histological transversal cuts in AT5, *athb5*, and WT plants. AT5 plants exhibited flattened petioles, less phloem area, shorter vascular bundles with a reduced area, and fewer xylem vessels than the WT control, whereas *athb5* mutant plants had the opposite characteristics (Figure 2C). Quantification of vascular length, xylem vessels number, and epidermis area indicated that AT5 plants showed a reduction in these parameters, while *athb5* mutants had an enhancement, both compared with the WT.

Because carbohydrate transport was affected in AT5 plants and whole-partitioning requires the specific activity of membrane sugar transporters, expression levels of different *SUC* and *SWEET* genes were evaluated. Transcript levels of *AtSUC1-9* and *AtSWEET11*-17 did not show significant changes between genotypes (Supplementary Figure S4), indicating that these genes were not transcriptionally affected by AtHB5.

Considering the latter result, we wondered if the differences observed in sugar accumulation could be the consequence of callose deposition in the phloem. Such deposits usually obstruct the correct transport from source to sink tissues (van Bel, 2003; Julius *et al*., 2018). Leaves of plants from the three genotypes (mutant, overexpressor, and WT) were stained with aniline blue. Notably, callose was detected only in overexpressor plants, suggesting that this could be the reason for starch accumulation (Figure 2D). Further, to know if AtHB5 could be regulating callose synthase gene expression, we evaluated transcript levels of *GSL3, GSL83*, and *GSL12* on *athb5*, AT5, and WT plants; none of them exhibited significant changes (Supplementary Figure S5).

### Starch metabolism is altered in AtHB5 mutant and overexpressor plants

Regarding starch accumulation observed at the end of the day in AT5 plants, we wondered whether the synthesis or degradation of this reserve carbohydrate was affected. Transcripts levels of key enzymes participating in starch metabolism were tested in 3-week old leaves four hours after the beginning of the day (10 am). Among them, SS1 (SUCROSE SYNTHASE 1) participates in the amylopectin synthesis pathway, SS5 (SUCROSE SYNTHASE 4) promotes starch granule initiation, and GBSS is a granule-bound starch synthase (Szydlowski *et al*., 2011; Abt *et al*., 2020). Both *SS1* and *SS4* levels were upregulated in AT5 but unchanged in *athb5* mutants compared to WT. Oppositely, *GBSS* was induced in *athb5* mutants and repressed only in one of the overexpressor lines compared to WT (Figure 3).

**Figure 3.**
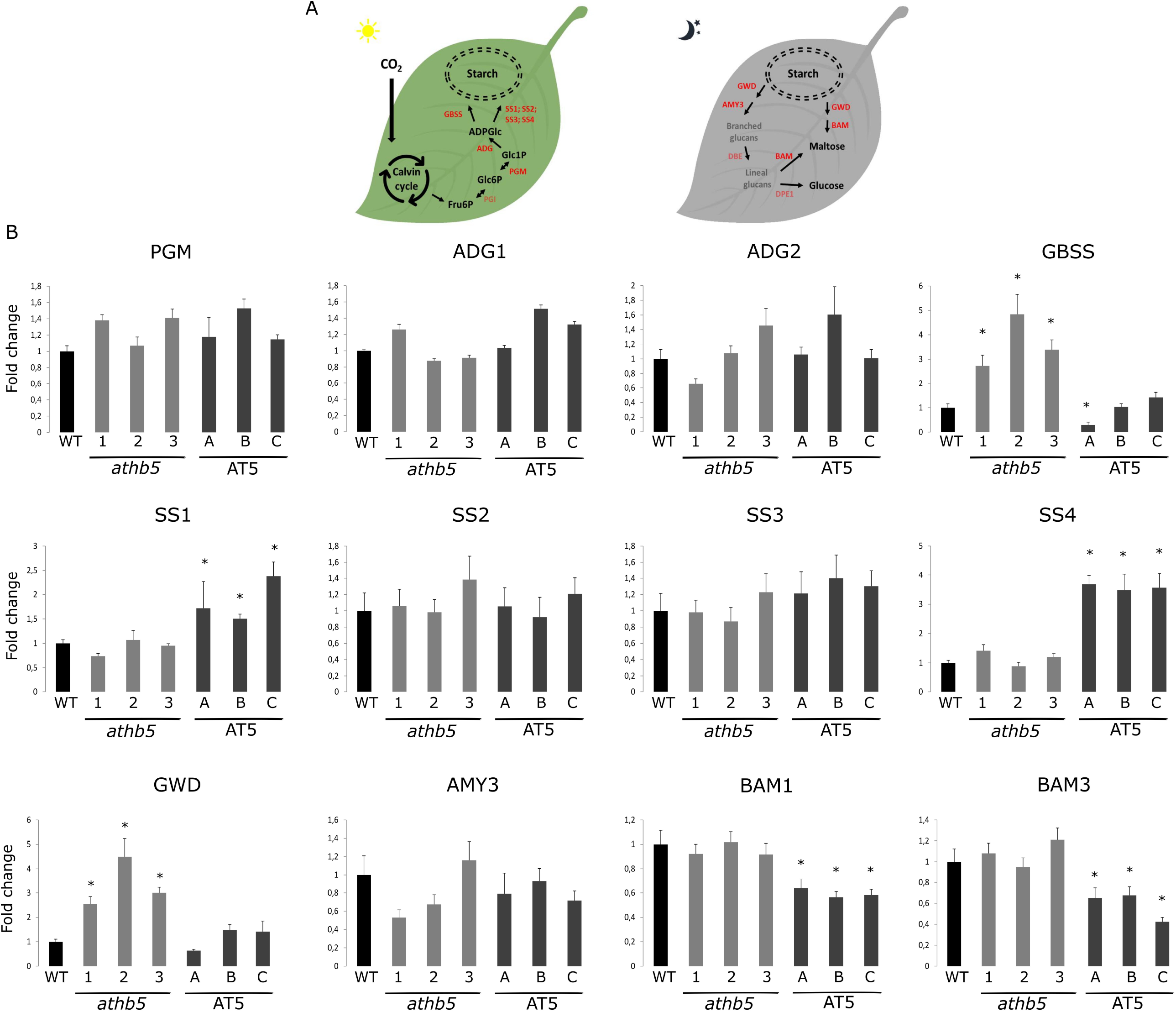
Transcript levels of genes involved in starch synthesis and degradation are differentially modulated in *athb5* and AT5 plants. (A) Illustrative picture indicating the function of genes encoding enzymes participating in starch synthesis and degradation. (B) *SS1, SS4, GBSS, GWD, BAM1*, and *BAM3* transcript levels were quantified by RT-qPCR in *athb5* mutant, AT5 overexpressor, and WT plants. The values were normalized with the one obtained in WT. The experiment was performed with four biological replicates, obtained by pooling tissue from four individual plants and tested by duplicate The bars represent SEM. Differences were considered significant and indicated with asterisks when the P-value was <0.05 (Student’s t-test).

ADP-GLUCOSE PYROPHOSPHORYLASE is composed of small and large subunits. The small one, encoded by *ADG1*, exerts the catalytic activity, while *ADG2* encodes the large one that catalyzes a rate-limiting step of starch biosynthesis. PGM (PHOSPHOGLUCOMUTASE) catalyzes the reversible conversion of glucose 6-phosphate (Glc6P) to glucose 1-phosphate (Glc1P). None of the transcript levels corresponding to these enzymes showed significant changes between genotypes (Figure 3). Regarding starch degradation, GWD (GLUCAN WATER DIKINASE), PWD (PHOSPHOGLUCAN WATER DIKINASE), BAM proteins (β-AMYLASES), and AMY3 (ENDOAMYLASE) play central roles. *GWD* was induced in *athb5* mutant plants but did not change in AT5 ones, both compared to WT. Both *BAM1* and *BAM3* were repressed in AT5 plants and did not change in *athb5* mutants (Figure 3). These results suggested starch synthesis and degradation are induced and repressed in AT5 and *athb5* plants, respectively.

### Source-to-sink carbon portioning is affected by AtHB5 levels impacting seed yield

To better understand the impact of AtHB5 in carbon partitioning, we evaluated physiological parameters in mutants and overexpressors along the plant life cycle. The analysis was carried out in two developmental stages: the juvenile stage was defined when the stem reached 10 cm height and the mature stage when it had 30 cm. The differences in rosette leaf dry-weight between genotypes were only appreciated in one AT5 line at the juvenile stage, whereas the other overexpressors showed solely a tendency (not statistically significant) to have scarcer biomass than controls (Figure 4A). Regarding stem biomass, the differences were significant; *athb5* mutants had more biomass at both stages, while overexpressors showed the opposite phenotype at the mature stage and a similar tendency at the juvenile stage (Figure 4B).

**Figure 4.**
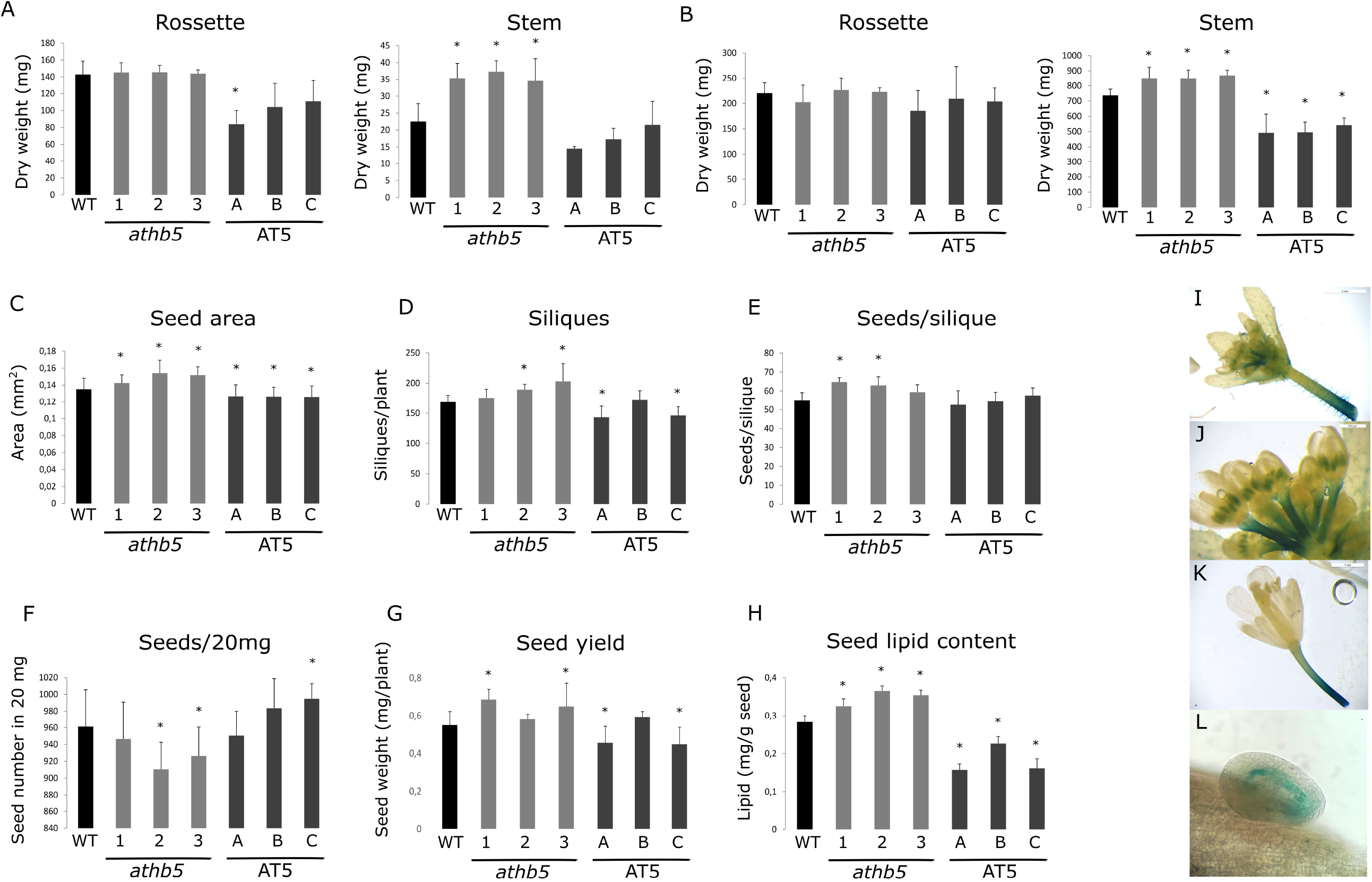
AtHB5 affects stem morphology, seed yield and content. The dry weight of rosette and stem biomass of 10 cm (A) and 30 cm (B) stem-length *athb5* mutant, AT5 overexpressor, and WT plants. (C) and (D) seed area (mm^2^) and lipid content (mg lipids/g seeds) of the same plants. Histochemical detection of GUS enzymatic activity in *ProAtHB5:GUS* immature flowers (E, F), pedicel of mature flowers (G), and seeds (H). The values were normalized with the one obtained in WT. Dry weight evaluation was performed with four biological replicates per genotype. Lipid determination was repeated twice with four biological replicates randomized from four individual plants per genotype. Bars represent SEM. Differences were considered significant and indicated with asterisks when the P-value was <0.05 (Student’s t-test).

Regarding the reproductive stage, *athb5* mutants had more siliques per plant (Figure 4C) and seeds per silique than WT plants (Figure 4D), while AT5 plants exhibited the opposite phenotype. Moreover, *athb5* mutant seeds showed an enlarged seed area (Figure 4E) and a smaller seed number each 20 mg, indicating that seeds were heavier than those of the WT (Figure 4F). In parallel, AT5 plants showed the opposite phenotype. As a result, *athb5* mutant plants yielded more than the WT, indicating that this TF is a negative modulator of seed production (Figure 4G). Stating the hypothesis that carbohydrate transport from source to sink tissues diminished in AT5 plants and this fact must impact seed quality, we analyzed the lipid content in seeds. The results indicated that lipid content is depressed in overexpressors and improved in mutants, both in the three independent lines and in agreement with the hypothesis (Figure 4E).

To confirm that the differential parameters (seed area, lipid content, seed yield) are closely related to *AtHB5* levels, we evaluated its expression pattern in reproductive tissues and organs. GUS staining was evident in the peduncle and anthers of immature flowers (Figures 4I, 4J), and in a more advanced developmental stage, in the peduncle of flowers and seed endosperm (Figures 4K, 4L).

### AtHB5 inhibits the transport of sucrose in stems

The leaf central vein transports sucrose through the petiole; then, the carbohydrate is mobilized by the stem vascular bundles to sink tissues. Given the enhanced stem biomass and the increased seed yield in *athb5* mutants, we wondered whether sucrose transport was arrested in stems. To better understand this scenario, the branches of adult plants were cut-off, leaving plants only with one main stem. This experiment forced the transport from rosette leaves to flowers via the main stem phloem, avoiding sucrose supply from the cauline leaves (Figure 5A). Sucrose was quantified in stems 24 h after cutting the branches, and the results indicated that the stems of *athb5* mutants had more sucrose than those of controls, whereas overexpressors showed the opposite behavior (Figure 5B). This result suggested that mutants have more transported sucrose, explaining grain yield and that AtHB5 is a transport inhibitor. When the life cycle ended, we evaluated the length of siliques and the seed weight. The cut of branches affected silique development producing shorter or not well-developed ones. This negative effect depended on the genotype; i.e., mutants were less penalized than controls and overexpressors (Figure 5C). The ratio between defective and total siliques was 13-33 % in mutants, whereas this percentage reached 67-74 % for overexpressors and around 50 % for the WT control. Moreover, seed yield in debranched plants showed a stronger genotype dependence than in intact plants; *athb5* mutants produced more and AT5 fewer seeds than WT (Figure 5D).

**Figure 5.**
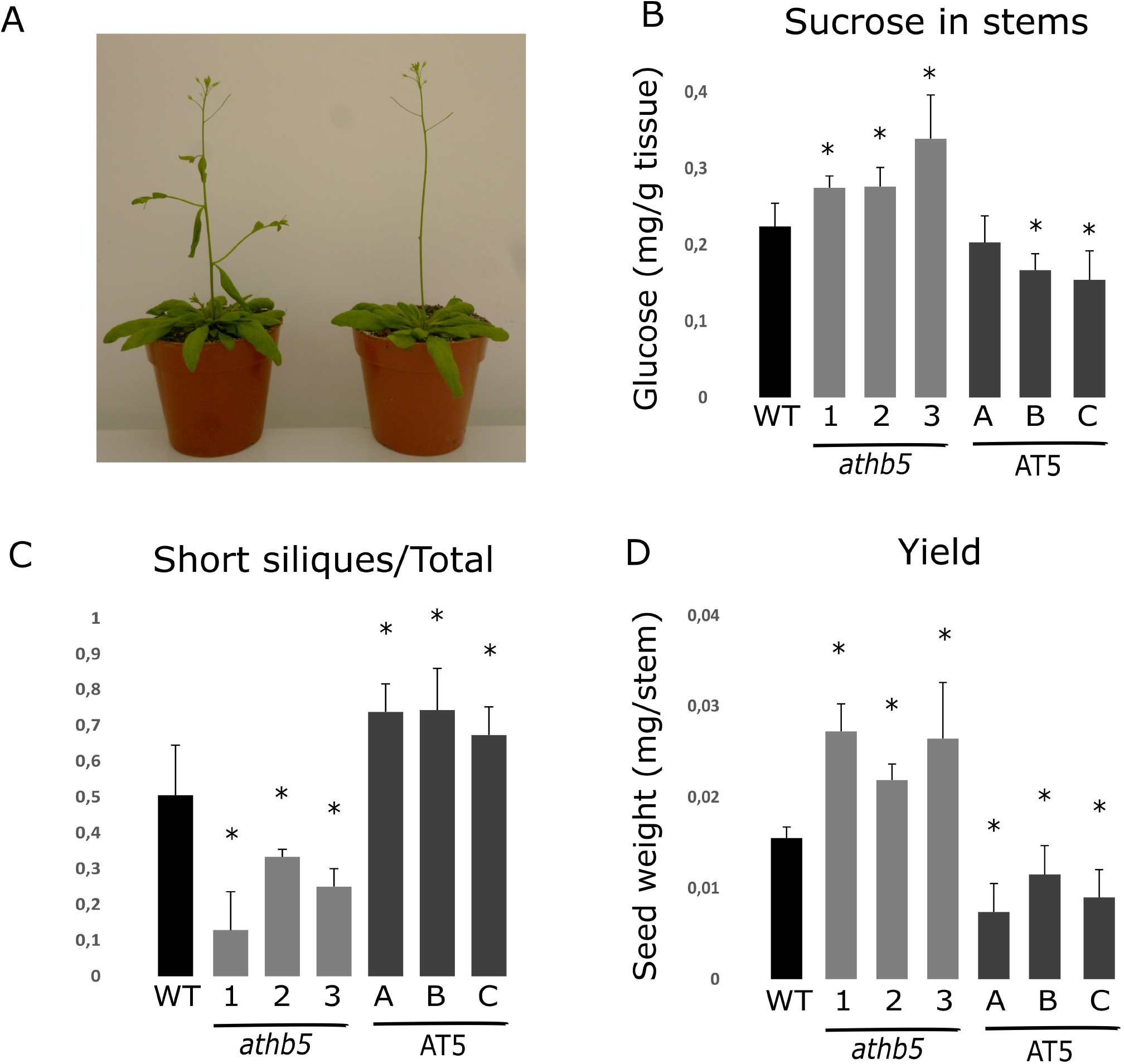
Forcing carbohydrate transport through the main stem shows the negative role of AtHB5 in source to sink carbon partitioning. (A) Illustrative photo of Arabidopsis plants before and after cutting off the branches. (B) Stem sucrose content in *athb5* mutant, AT5 overexpressor, and WT debranched plants, expressed as mg of glucose per g of fresh weight. (C) Short siliques over total siliques ratio in the same plants. (D) Seed yield (mg/individual stem). Experiments were performed with eight plants per genotype and repeated at least twice. Carbohydrate determination was repeated twice with four biological replicates randomized from eight individual plants per genotype Bars represent SEM. Differences were considered significant and indicated with asterisks when the P-values were <0.05 (Student’s t-test).

### Sucrose transport to roots is also affected by AtHB5

Roots are sink organs during the vegetative stage of development. To evaluate the putative role of AtHB5 in carbohydrate transport to roots, *athb5* mutant and AT5 plants were grown on MS-agar under normal conditions.

We analyzed the expression pattern using the *ProAtHB5:GUS* plants; staining was evident in hypocotyls and the upper zone of the primary root vascular tissue of five-day-old seedlings (Figure 6A). Regarding that *AtHB5* was expressed in roots, we analyzed the root phenotype of 12-day-old *athb5* mutants and AT5 overexpressors (Figures 6B and 6C). The latter exhibited shorter roots and fewer emerged lateral roots compared with controls, whereas a slight opposite phenotype was observed in the mutants. When plants grew in darkness, sucrose was limited. In six-day-old seedlings grown in MS medium supplemented, or not, with 3% sucrose or sorbitol as osmotic control, all the plants were able to use the sucrose and elongated their roots. However, *athb5* mutants were more successful than controls and overexpressors. In the plate supplemented with sorbitol, all the roots elongated but to a low extent, and no differences were detected between genotypes, discarding osmotic stress as the factor generating differential phenotypes in the plate supplemented with sucrose (Figure 6D).

**Figure 6.**
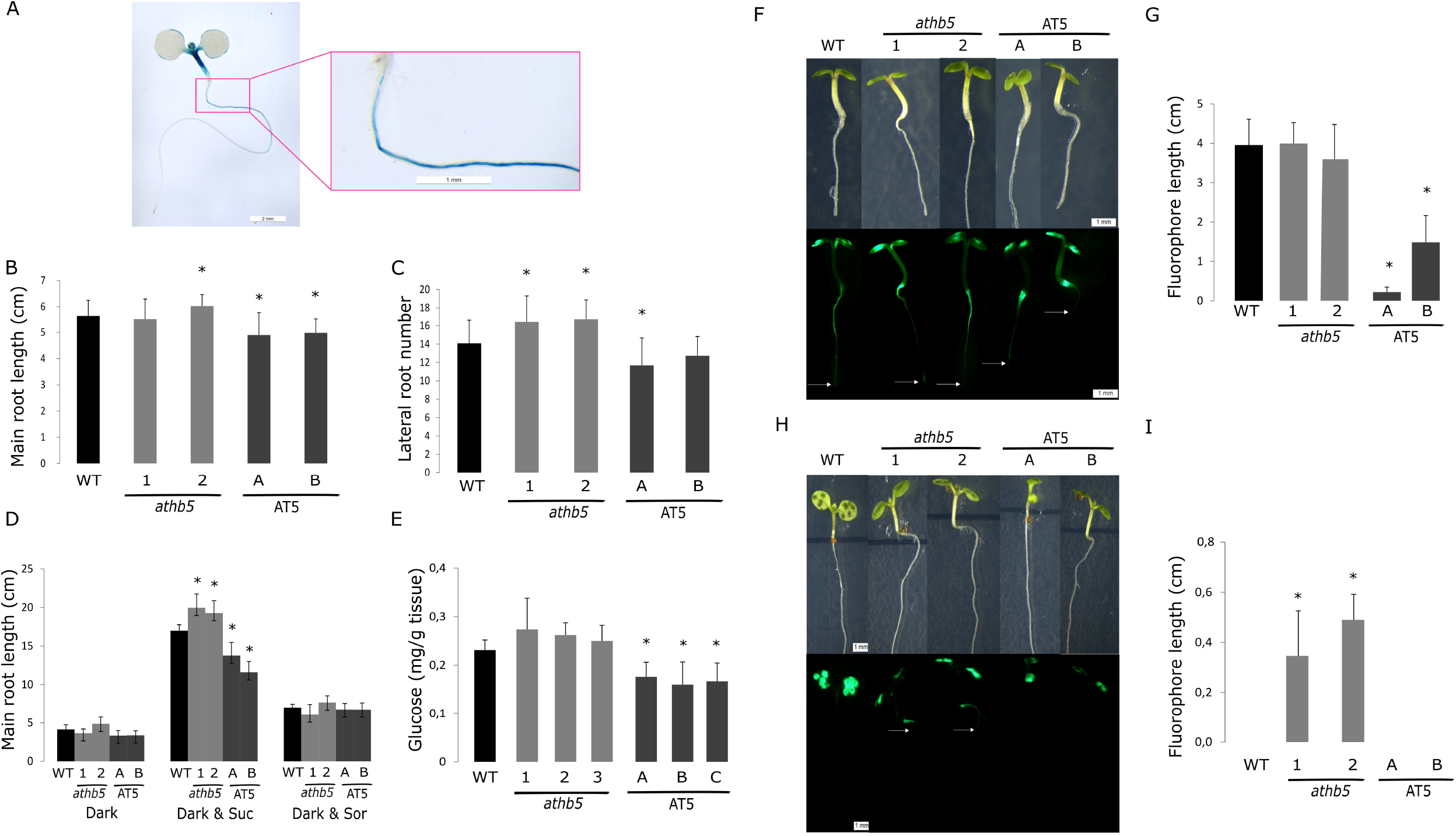
AtHB5 modulates root morphology and inhibits sucrose transport to roots. (A) Histochemical detection of GUS enzymatic activity 7-day-old *ProAtHB5:GUS* seedlings. (B) Primary root length and (C) number of lateral roots of 7-day-old *athb5* mutant, AT5 overexpressor, and WT plants seedlings grown under normal conditions. (D) Primary root length of seedlings grown in darkness and supplemented (or not) with sucrose (3%) or sorbitol. (E) Sucrose content (mg/g of fresh tissue) in the roots of the same genotypes, grown in normal conditions. (F) Upper panel, bright field of 7-day old seedlings (*athb5*, AT5, and WT). Lower panel: CFDA probe transported in the same plants, visualized with a fluorescence microscope to detect the probe after 10 minutes of CFDA application in cotyledons. (G) Quantification with ImageJ of CFDA distance transported through the roots (cm). (H) Upper panel, bright field of 7-day old seedlings (*athb5*, AT5, and WT). Lower panel: CFDA probe visualized with a fluorescence microscope after 3 minutes of CFDA application in cotyledons. (I) Quantification with ImageJ of CFDA distance transported through the roots (cm). Quantification of root length was repeated twice and performed with 20 seedlings per genotype and condition. CFDA experiments were performed with at least four biological replicates per genotype and repeated twice. Bars represent SEM. Differences were considered significant and indicated with asterisks when the P-values were <0.05 (Student’s t-test). White scale bars represent 1 mm. White arrows indicate the distance reach by the fluorescent probe.

To evaluate sucrose content in roots receiving this carbohydrate only from the source tissue, 10-day-old seedlings were harvested and evaluated. AT5 plants exhibited lower sucrose content than controls, whereas the differences in the mutants were not statistically significant (Figure 6E). This observation suggested a problem in carbohydrate transport through the phloem in the overexpressors. To confirm this, we used the fluorescent probe CFDA that can be loaded into source leaves, monitoring the phloem streaming and unloading in the sink tissues by fluorescence. Plants from the three genotypes were grown in vertical MS plates without sucrose supplementation for four days. CFDA was applied to the cotyledons and visualized 10 min after (Figure 6F). All the plants showed fluorescence in the root phloem, indicating a successful dye loading, and the followed path was quantified using ImageJ. The results indicated that AT5 plants showed a shorter probe route compared with the WT or *athb5* mutants (Fig 6G). Shorter times were needed to visualize differences between athb5 mutant and WT plants. CFDA was applied, visualized 3 min after, and fluorescence pathway length quantified (Figure 6H). The results showed that *athb5* mutants could transport CFDA faster than WT.

## Discussion

Transcription factors belonging to the HD-Zip I family were associated with plant development and in response to abiotic stress factors (Perotti *et al*., 2017). Among HD-Zip I TFs, only *AtHB5* was repressed during the passage from immature to mature stems (Ko *et al*., 2004). Despite all the background information about AtHB5 (Stamm *et al*., 2012, 2017), its role in stem maturation was unknown.

The analysis of *athb5* mutants and *AtHB5* overexpressors indicated significant differences in stem growth, specifically in stem width. Wider stems were previously associated with enhanced yield (Cabello and Chan, 2019) and constitute a desirable agronomic trait. In the same sense, vascular bundles are crucial for water and nutrient transport (Ding *et al*., 2020). Notably, there are no reports on Arabidopsis HD-Zip I mutants exhibiting improved characteristics. In contrast, there are several works in which the over or ectopic expression of these genes conferred potential beneficial traits, such as increased tolerance to abiotic stress factors, in many plant species (Manavella *et al*., 2006; Cabello *et al*., 2012; Cabello *et al*., 2016; Y. Zhao *et al*., 2014; Basso *et al*., 2021; S. Zhao *et al*., 2021; Raineri *et al*., 2022). Moreover, several constructs able to overexpress HD-Zip I members and others to knock-down these genes were protected with patents (Arce *et al*., 2008; Guo *et al*., 2011; Sanz-Molinero *et al*., 2016). However, in other species like rice, *OsHOX22* mutant plants were described as drought tolerant (Zhang *et al*., 2012). Phylogenetic trees indicated that OsHOX22 is the rice homolog of the Arabidopsis AtHB7 and AtHB12 transcription factors (Agalou *et al*., 2008).

Petiole and leaf morphology were associated with starch storage ability and water status (Raines and Paul, 2006; Shen *et al*., 2019; Falchi *et al*., 2020). *AtHB5* mutant and overexpressors exhibited an altered opposite morphology considering petiole and vascular bundle area and xylem and phloem vessels, indicating a role of this gene in the determination of conductive tissue morphology. Supporting a role of AtHB5 in carbohydrate transport, the overexpressors showed callose deposition. Such deposition constitutes one of the mechanisms affecting carbohydrate transport through the phloem. The pores located at the ends of the sieve vary in size and impact the flow of phloem sap throughout the plant (Barratt *et al*., 2011; Xie *et al*., 2011). Hence, the enhanced callose deposition observed in AT5 plants could explain, at least in part, the reduced carbohydrate transport in such plants, supporting a negative role of sugar transport for AtHB5. Maize carbohydrate partitioning defective1 (*cpd1*) was identified as a semi-dominant mutant exhibiting hyperaccumulation of starch and soluble sugars in leaves. This accumulation was due to an inhibited sucrose export than could be explained by ectopic callose deposits in the phloem of mutant leaves. As a consequence, *cpd1* mutant plants showed reduced height and yield (Julius *et al*., 2018), similar to the phenotype observed in AT5 plants.

Regarding the regulation of genes involved in starch synthesis, *SS1* and *SS4* were induced in AT5 plants. The encoded enzymes participate in amylopectin synthesis. SS1 generates short-chain glucans (Hizukuri *et al*., 1986), whereas SS4 is not required for the normal amylopectin structure formation and plays a role in granule initiation (Roldán *et al*., 2007). The induction of these genes in AT5 plants would suggest that AtHB5 positively regulates the generation of short-chain glucans of amylopectin and the formation of starch granules. On the other side, amylose synthesis is strictly dependent on GBSS and does not require any of the four sucrose synthases. *GBSS* expression is induced in *athb5* mutants, indicating that AtHB5 could negatively regulate amylose synthesis.

Considering genes involved in starch degradation, *GWD* was up-regulated in *athb5* mutants, while *BAM1* and *BAM3* were down-regulated in AT5 plants, both compared with the WT. *GWD1* and *PWD* encode water dikinases that phosphorylate at the glucose C6 and C3 positions, respectively (Ritte *et al*., 2002; Baunsgaard *et al*., 2005; Kötting *et al*., 2005). BAM3 and BAM1 are two of the nine Arabidopsis β-amylases that hydrolysate the outer starch chains releasing maltose (Fulton *et al*., 2008). These results suggested that AtHB5 could be repressing the first step of starch degradation and slowing down the release of maltose. Approximately 80% of fixed C is exported to sink tissues, and the regulation of carbon partitioning is vital for plant growth and development. Plants exhibiting impaired starch synthesis or degradation had reduced biomass in normal photoperiod growth conditions but not under continuous light or very long days (Caspar *et al*., 1985; Gibon *et al*., 2004, 2009). This is because fewer starch reserves negatively impact plant growth during the night (Wiese *et al*., 2007).

Plants with altered *AtHB5* levels exhibited significant changes in stem biomass; *athb5* mutants exhibited enhanced biomass, whereas overexpressors showed the opposite phenotype, indicating that AtHB5 negatively influences growth rate. In agreement, sucrose, glucose, and starch augmented in the mutants. All these differential traits impact seed yield and composition. Debranching the plants provoked the enhancement of this differential phenotype, supporting that long-distance transport is favored in *athb5* mutants compared with WT and AT5 plants. Studying the signals displayed by trehalose-6P in axillary bud outgrowth after decapitation, different carbohydrates in pea decapitated plants were evaluated (Fichtner *et al*., 2017). Trehalose-6P and sucrose significantly increased in stem samples, whereas glucose showed little changes. This experiment aimed to eliminate sucrose supply from leaves to determine whether the trehalose-6P signal depended on leaves as source organs. Defoliation was also used to explore the importance of shoot-derived nutrients for bud release (Mason *et al*., 2014).

Previous studies showed that root growth is influenced by direct root illumination and by glucose or sucrose supplementation (Silva-Navas *et al*., 2015, López-García *et al*., 2020, García-González *et al*., 2021a). Sucrose supplementation positively affected total root length in darkness but less than in light without the carbohydrate addition. Furthermore, the replacement of sucrose with glucose was not efficient in counteracting darkness, indicating that the carbon source had a crucial impact on root growth (García-González *et al*., 2021b). Here we demonstrated that in darkness, *athb5* mutants elongated better the primary root in sucrose-supplemented than the WT, whereas AT5 plants exhibited shorter roots than WT, indicating that *athb5* roots were more able to efficiently uptake sucrose from the medium than WT or AT5 plants.

In seedlings, the root is a major sink organ and depends on sucrose transported from cotyledons. Here we show that AT5 plants had less sucrose on roots under normal growth conditions and that the CFDA probe moved more quickly in *athb5* mutant plants, further supporting the negative role of AtHB5 in sucrose transport.

In summary, the results indicated that AtHB5 is a negative modulator of carbon partitioning from source to sink tissues in different developmental stages, modulating morphological, physiological, and biochemical traits impacting the performance of the whole plant.

## EXPERIMENTAL PROCEDURES

### Plant material and growth conditions

*Arabidopsis thaliana* plants (Col-0 ecotype) were grown on Klasmann Substrat Nº 1 compost (Klasmann-Deilmann GmbH, Germany) in a growth chamber at 22-24 °C under long-day (16/8 h light/dark cycles) conditions, with a light intensity of approximately 120 µmol m^-2^ s^-1^ in 8 × 7 cm pots. Four plants were planted per pot, unless stated differently.

For root phenotyping: seeds (WT, mutant, and overexpressor plants) were surface sterilized and placed at 1 cm from the top of square Petri dishes (12 × 12 cm) for 3 days at 4 °C before placing the dishes in the growth chamber at 22-24 °C under long-day (16/8 light/dark cycles) with a light intensity of approximately 70 µmol m^-2^ s^-1^. The growing medium was Murashige-Skoog supplemented with vitamins (MS, PhytoTechnology Laboratories™).

Arabidopsis mutant lines *athb5*-1, -2, and -3 (SALK_015070c, SALK_134509 and SALK_078606) were obtained from the ABRC

### Genetic constructs

#### 35S:AtHB5

*AtHB5* cDNA was obtained by RT-PCR of RNA isolated from Arabidopsis Col-0 14-day-old seedlings and cloned in the pGEM T-easy plasmid. Then, the cDNA was cut with *BamH1/KpnI* and inserted directly in the pBI121 previously restricted with the same enzymes, replacing the *GUS* reporter gene. Specific oligonucleotides were described in Supplementary Table S1.

#### PromAtHB5:GUS

A 1531 bp segment upstream of the transcription start site was isolated from Arabidopsis genomic DNA with specific oligonucleotides (Supplementary Table S1) and cloned into the pGEM®-T Easy (Promega). After that, the fragment was restricted with *EcoR*I and subcloned into the pBluescript SK-vector. After checking the correct orientation, the DNA was treated with *Hind*III and *BamH*I and finally inserted into the pBI101.3 vector. The correct cloning was verified by sequencing. The final construct, able to express GUS was used under the control of the *AtHB5* promoter, and was used to transform plants. The constructs were used to transform *E. coli* DH5α cells, and DNA prepared for sequence determination (Macrogen, Korea). Once checked, it was used to transform *Agrobacterium tumefaciens* cells (LBA4404).

### Histology and microscopy

Arabidopsis tissues (first internodes and petioles) were harvested and fixed as previously described (Cabello and Chan, 2019).

Safranin-Fast green staining was used for xylem/phloem differential detection. After removing paraffin with 100 % xylene for 15 min at room temperature, sections were imbibed in ethanol (100%, 96% and 90%), then the slices were transferred to Safranin (in ethanol 80%) for at least 4 h. After that, slices were put in ethanol (90%, 96% and 100%) to finally be imbibed on Fast-green dye (in ethanol 100%) during seconds and washed in ethanol 100% to remove excess of dye. Slices were mounted on Canadian balsam.

For lignin staining, paraffin from slices was removed using xylene 100 %. Then they were dehydrated in ethanol 100% and treated with Fluoroglucinol-HCl during 30 minutes. After that, the exceeding dye was removed and rapidly mounted on Canadian balsam. Lignin was visualized immediately under optical microscope.

For GUS staining, paraffin from slices was removed using xylem 100%. Then they were dehydrated in ethanol 100% and rapidly mounted on Canadian balsam.

When the intensity of the expression in certain tissues was difficult to visualize on 10 µM cross-sections, photographs were initially taken from whole paraffin inclusions. For callose staining, Arabidopsis leaves were harvested, cleared by immersing them in 70 % ethanol, and stained with aniline blue for callose detection as described Hauck *et al*., (2003). Then, they were examined with a Leica fluorescence microscope.

For determining different petiole and stem area, histological sections from at least four biological replicates were photographed and then measured using the ImageJ software (NIH, Maryland, United States).

### Histochemical GUS staining

In situ assays of GUS activity were performed as described by Jefferson *et al*., (1987). Whole plants were immersed in a 1 mM 5-bromo-4-chloro-3-indolyl-b-glucuronic acid solution in 100 mM sodium phosphate pH 7.0 and 0.1 % Triton X-100 and, after applying vacuum for 5 min three times, they were incubated at 37 °C overnight. Chlorophyll was cleared from green plant tissues by immersing them in 70 % ethanol. Paraffin inclusions were performed after chlorophyll was totally cleared.

### Plant phenotyping

Plant architecture parameters were scored manually or with the aid of a ruler or gauge. All the experiments were performed with four to eight plants per genotype and per treatment and repeated at least 3 times.

To determine the dry weight, fresh tissue was placed at 37°C for four to six days, until the weight was constant.

For determining petiole and stem parameters, *athb5* mutant, AT5 overexpressor, and WT plants were grown in individual pots, and petioles from rosette leaves or first internode sections were cut and mounted on paraffin blocks as previously described.

For determining seed parameters, plants were harvested at maturity, when siliques werefully ripped. Twenty grams of seeds were weighted and then counted (N = 10). For determining area (N = 20), dried seeds were photographed and then measured using the ImageJ software (NIH, Maryland, United States).

Root phenotype was determined in seedlings grown on MS plates and quantification was performed with at least 20 individuals per genotype and condition. The experiment was repeated twice.

### Soluble sugars, starch and lipid content

For carbohydrate measurements, plant material (50-80 mg each sample) was frozen and ground to powder in liquid nitrogen and extracted with a solution containing 5,4 mM phosphate buffer pH 7,5; 0,1 mM EDTA, 62,5 % metanol, and 26,8 % chloroform. The samples were incubated on ice for 20 minutes and after that, 300 µl of water were added, mixed and centrifuged for 5 min at 13,000 rpm. The supernatant was dried overnight at 40 ºC and used to quantify sucrose and glucose. After resuspension, a fraction was used to determine glucose and another fraction was treated with invertase to determine sucrose. For starch extraction, the pellet was resuspendend in 1 mL of ethanol and centrifuged at 13000 rpm for 5 minutes, dried at 70 ºC, and finally resuspended with 250 µl of 0,1 N NaOH. Once a solution was obtained, 75 µl 0,1 N AcH (pH 5,1) were added. Each sample (50 µl) was treated with amiloglucosidase overnight at 37 ºC. Glucose was measured using a kit containing glucose-oxidase (GOD), peroxidase (POD), 4-aminofenazone (4-AF), phosphate buffer (pH 7), and 4-hydroxibenzoate. The DO was measured at 505 nm and glucose concentration was calculated using a glucose standard solution.

The extraction and quantification of lipids by gravimetry was based on the technique described by Siloto *et al*. (2006) and adapted for small volumes. Around 25 mg of seeds were ground and incubated with 400 µl of isopropanol for 10 min, at 65 °C, and then air-dried. Lipids were obtained after three extractions of methanol, chloroform and water biphasic solution (methanol:CHCl_3_:H_2_O). The first extraction was performed with 1.0 mL of methanol:CHCl_3_:H_2_O (2:2:1.8 [v/v]). The second and third extractions were performed with 1.0 mL of methanol:CHCl_3_:H_2_O (1:2:0.8 [v/v]). These three extractions were separated by centrifugations at 6700 g during 5 min. The lipid fractions were collected and the solvents were completely evaporated. All the fractions were collected in a unique previously weighted microtube (W1), and after drying of the sample, it was weighted again (W2). Lipids were expressed as W2 − W1/mg tissue. These experiments were repeated at least twice. Samples were obtained by pooling tissue from four to eight individual plants per genotype.

### RNA isolation and expression analyses by RT-qPCR

Total RNA for real-time RT-PCR was isolated from rosette leaves of 25-day-old plants using Trizol® reagent (Invitrogen, Carlsbad, CA, USA) according to the manufacturer’s instructions. Total RNA (1 μg) was reverse-transcribed using oligo(dT)_18_ and M-MLV reverse transcriptase II (Promega, Fitchburg, WI, USA).

Quantitative real-time PCR (qPCR) was performed using an Mx3000P Multiplex qPCR system (Stratagene, La Jolla, CA, USA) as described before (Cabello *et al*., 2016) and using the primers listed in Table S1. Transcript levels were normalized by applying the ΔΔCt method. Actin transcripts (*ACTIN2* and *ACTIN8*) were used as internal standards. Four biological replicates, obtained by pooling tissue from 4 individual plants and tested by duplicate, were used to calculate the standard deviation.

### CFDA Application

CFDA (6 mg/mL) was prepared in acetone and kept at -80°C. A 1:20 dilution in sterile water was used for application. A cotyledon was grazed by fine tweezers to allow 1 μL CFDA to penetrate. All samples were excited by a 488-nm laser. Fluorescence 500 to 566 nm was monitored. Roots were photographed after 5 or 10 minutes of application using an adapted Nikon camera. Quantification o CFDA transport was performed using ImageJ software.

### Accession codes

SS1 (AT5G24300); SS2 (At3g01180); SS3 (At1g11720); SS4 (AT4G18240); GBSS (AT1G32900); ADG1 (At5g48300); ADG2 (At5g19220); PGM (AT5G51820); GWD (AT1G10760); BAM1 (AT3G23920); BAM3 (AT4G17090); AMY3 (AT1G69830).

## Acknowledgements

This work was supported by Agencia Nacional de Promoción Científica y Tecnológica (PICT 2017 0305 and PICT 2019 01916 to RLC, PICT 2015 1286, and CAI+D 2016 to JVC) and CONICET. LR and VM are CONICET Ph.D. Fellows, and CZ was undergraduate student. JVC and RLC are CONICET Career members.

## Conflict of interest

The authors declare no conflict of interest

## Authors’ contribution

Conceived and designed the experiments: JVC and RLC. Performed the experiments: LR, VM, CZ, and JVC. Analyzed the data: JVC and RLC. Wrote the paper: JVC and RLC.

## Legends to Supplementary Material

**Figure S1. *AtHB5* expression levels in AT5 plants**

**Figure S2. The length and width of the main stem differ between *athb5*, AT5, and WT plants**

**Figure S3. Starch, glucose, and sucrose concentration during day and night**

**Figure S4. Expression levels of sucrose transporters in AT5 and *athb5* plants**

**Figure S5. Expression levels of callose synthases genes**

**Table S1. Oligonucleotides used for cloning and RT-qPCR assays**

## Notes

### Competing Interest Statement

The authors have declared no competing interest.

